# Single-cell sequencing for the molecular identity involved in zinc therapy for spinal cord injury

**DOI:** 10.1101/2023.01.19.524759

**Authors:** Zuqiang Shi, Jiaquan Lin, Minghao Ge, Hengshuo Hu, Liang Mao, Miaomiao Tian, He Tian, Xifan Mei

## Abstract

Spinal cord injury (SCI) is a severe traumatic neurological condition often caused by car accidents, violent impacts, or unintentional falls and can have devastating consequences for the patient. SCI include primary and secondary injuries. Followed by primary injury, and secondary injuries including inflammatory responses and oxidative stress, which exacerbate the disease process of SCI. Therefore, the treatment of SCI, especially the suppression of secondary injuries, is a major focus of attention in the field of neuroscience. In previous studies, we have demonstrated that zinc therapy can exert neuroprotective effects in mice after acute SCI, including reducing the inflammatory response in the central nervous system, decreasing the neuronal apoptosis and downregulating oxidative stress at the region of injury. However, little is known about how zinc therapy systematically alleviates SCI. Here, we have systematically analysed and mapped the single-cell atlas of the spinal cord in mice with SCI treated by zinc therapy, which provides a systematic analysis of the transcriptome of individual cells in the spinal cord. Among the results, we found that zinc therapy-induced alterations in the IL-17 inflammatory pathway and produced immune heterogeneity in microglia which are the inherent immune cells in the central nervous system. By cell subpopulations clustering analysis, we defined seven microglia subpopulations in zinc-therapy spinal cord tissue and identified the presence of novel VEGFA+ microglia. This finding presents that the microglia subpopulations affected by zinc therapy may have the potential to promote angiogenesis, which is a potential mechanism for the treatment of SCI. In conclusion, this study maps/reveals the transcriptomic of zinc therapy for the treatment of acute SCI with unprecedented resolution and provides a molecular basis for the clinical application.

## 1 Introduction

Spinal cord injury (SCI) is a serious traumatic neurological condition which often results in permanent partial or complete loss of sensory, motor and neuroautonomic functions [1]. SCI is divided into primary injuries, which usually result from mechanical damage, and secondary injuries, which are often accompanied by changes in tissue structure and the development of neuroinflammation[2,3]. SCI not only cause great emotional distress and inconvenience to the patient but also place a heavy burden on the family[4]. Therefore, the treatment of SCI is one of the concerns in the field of neuroscience.

Zinc, as a trace element, is often involved in biological behaviours such as proliferation promotion, anti-ageing, antioxidant and autophagy promotion in the free ionic state in living organisms[5–9]. When zinc gluconate is ingested by the body, zinc ions are absorbed into the blood via the stomach and are regulated by different zinc ion transport proteins[10,11]. In the central system, for example, zinc ions are mainly regulated by the zinc ion transport proteins Znt3 and ZIP8[10,11]. As zinc ions are involved in the normal development and daily function of spinal cord tissue, they are essential for the maintenance of normal physiological function of the central nervous system[12]. In previous studies, zinc ions were also found to be used in the treatment of SCI[13]. Welling et al. reported that zinc ion deficiency reduced the function of immune cells in the CNS, and in a follow-up study, they also found that zinc ions inhibited interleukin (IL)-1β expression in a dose-dependent manner, suggesting a potential anti-inflammatory effect of zinc treatment[14,15]. Meanwhile, in our previous study, zinc ions could exert neuroprotective effects after SCI by increasing the expression of brain-derived neurotrophic factor and granulocyte colony-stimulating factor, thereby promoting functional recovery after SCI[5,16]. Although the neuroprotective effects of zinc ions after SCI have been well demonstrated, the anti-inflammatory effects in the acute phase of SCI need to be further investigated.

Single-cell sequencing is a technique that allows the function of cells in normal physiological activities and pathological processes to be explored[17]. This technique has been used to discover cell types and their developmental trajectories in several organs, including the kidney and lung[18–20]. Many researchers have also used the technique to probe the function of cells in the central nervous system[21–23]. Similarly, to investigate the neuroprotective effects of zinc therapy and its anti-inflammatory effects in acute spinal cord injury, we used single-cell sequencing to try to elucidate the molecular mechanisms of zinc therapy at the cellular level. In this study, we sequenced single-cell RNA from 30,078 cells from 20 mice and mapped the transcriptomic profile with unprecedented resolution. By unbiased analysis, we annotated 18 cell clusters and 11 cell types, including microglia, neuronal cells and other common cell populations. Subsequently, we decoded the heterogeneity and stratification status of microglia after zinc treatment, further identifying seven clusters of microglia subpopulations and classifying them into five cell states, including the activated proliferative state (Mki67+) and the remaining four states (Ms4a7+, Ifit3+, Vegfa+ and Ppia+). Here, we systematically present the intercellular communication and signalling pathways involved in the environmental regulation of zinc treatment, particularly between microglia and the rest of the cells. We then established animal models and measured CD31 expression levels in tissue samples to reveal that zinc therapy can promote revascularisation at the site of injury, which is a potential mechanism for functional recovery from spinal cord injury. Thus, the findings of this study may provide a more comprehensive molecular basis for the clinical application of zinc ions in the treatment of spinal cord injury.

## 2. Methods

### 2.1 Animals

For the experiment, we used 66 (33 male and 33 female) eight-week-old C57BL/6J mice weighing 20–25 g in accordance with the National Council for Laboratory Animal Research’s “Guide for the Protection and Use of Laboratory Animals” (1996) and the “Guide for the Protection and Use of Animals” developed by the Animal Protection and Use Committee of Jinzhou Medical University. Before the experiment, the mice were housed at 22–24 °C in 12/12 h day/light cycles, with free access to food and water.

### 2.2 SCI model and zinc administration

The mice were randomly divided into sham, SCI + vehicle, and SCI + zinc groups. We used Allen’s method to build a spinal cord contusion model[24]. Briefly, the mice were first anaesthetized with isoflurane. After complete anaesthesia, the laminas were removed at the T9–T10 level without damaging the dura mater. The impactor (Cat#Model III, RWD Life Sciences, USA; 2-mm diameter, 12.5 g, 20-mm height) was dropped on the T9–T10 spinal cord surface, causing moderate spinal cord contusions. The wound was sutured and disinfected with iodophor, and the mice were placed at a steady temperature and in a well-lit environment. The sham group was operated on in the same manner except that it received a spinal cord contusion. The SCI + zinc group was administered with an intraperitoneal injection of zinc gluconate (30 mg/kg; Biotopped, Beijing, China) 2 h postoperatively and again 1 day later at the same dose until day 3. The SCI + vehicle group was assigned an equal amount of isotonic glucose solution simultaneously as a control.

### 2.3 Preparation of single-cell suspensions

Samples from the injured spinal cord on day 1 were isolated and quickly transported to the research facility. Each sample was subsequently minced on ice to <1 mm cubic pieces, followed by enzymatic digestion using type 2 collagenase (e.g., Sigma) with manual shaking every 5 min. Samples were then centrifuged at 300 RCF for 30 s at room temperature, and the supernatant was removed without disturbing the cell pellet. Next, 1× of phosphate-buffered saline (calcium- and magnesium-free) containing 0.04% w/v of bovine serum albumin (BSA; 400 μg/mL) was added and centrifuged at 300 RCF for 5 min. The cell suspension was then resuspended with 1 mL of erythrocyte lysis buffer and placed at 4 °C for 10 min. Subsequently, the sample was resuspended with 1 mL of 1× phosphate-buffered saline containing 0.04% w/v BSA (400 μg/mL) and filtered through a Scienceware Flowmi 40 μm cell filter. Finally, the cell concentration and viability were determined using toluidine blue staining and hemocytometry.

### 2.4 10× Genomics scRNA-seq

To interpret the multiplexed cell barcodes, we used 10× Genomics Cell Ranger software (version 5.0.0) and mapped the reads to the genome and transcriptome using STAR aligner. As required, we down-sampled the sample reads to generate normalized aggregated data across samples and matrices of gene counts to cells. We used the R package, with Seurat (version 3.1.1), to process unique molecular identifiers (UMIs) to count the matrices[25]. We assumed Gaussian distribution of the UMI/gene counts/cell and used it as a criterion to filter cells with UMI/gene counts exceeding +/- 2 times the standard deviation of the mean for possible multi-point capture and removal of low-quality cells. After analyzing the cellular distribution of the mitochondrial gene expression in proportion to its presentation, we discarded low-quality cells belonging to mitochondrial genes with a threshold >10%. Simultaneously, we used the DoubletFinder package to identify potential doublets. Individual cells from the spinal cord were included in downstream analyses after applying these quality control criteria. We then used the data normalization function in the Seurat package to normalize the library size and, in turn, obtain normalized counts. In particular, we normalized the gene expression measurements of each cell by the total expression using the global scale normalization method of lognormalizing, multiplied the results by a scale factor (default 10,000), and log-transformed the results. Macosko et al.’s method to determine the top variable genes in individual cells was applied[26]. We then used the FindVariableGenes function in the Seurat package to select the most variable genes and performed graph-based clustering based on the gene expression profile of the cells using the FindClusters function in the package. We used the RunTSNE function in the package and the 2D student t-distributed stochastic neighbor embedding (t-SNE) algorithm for cell visualization. The FindAllMarkers function (test.use=bimod) in the Seurat package was then used to identify the marker genes for each cell cluster. For each cell cluster, we identified the markers that were positive compared to the other cells. Finally, we used the R Package SingleR algorithm for unbiased cell type identification by scRNA-seq while referring to the transcriptional dataset “Human Primary Cell Atlas” to infer the cell of origin of each individual cell and perform cell-type identification[27].

We used the FindMarkers function in Seurat to identify differentially expressed genes. A p-value < 0.05 and |log2foldchange| > 0.58 were set as differential expression thresholds. Moreover, we analyzed gene oncology enrichment and KEGG pathway enrichment of differentially expressed genes using the hypergeometric distribution of R. Sequencing and bioinformatics analysis were performed by OE Biotech (Shanghai, China).

### 2.5 Pseudotime analysis

We used the Monocle2 package to determine the developmental pseudotime of the cells. First, we used the import CDS function to convert the Seurat object into a CellDataSet object. We then used the DifferenceGenetest function in the package to sort genes (p-value < 0.01) and assist in sorting cells on the pseudo-time track. We then used the reduceDimension position to perform a reduced dimensional clustering analysis and the default parameters of the orderCells function to perform trajectory inference. Finally, we used the plot_genes_in_pseudotime function to plot gene expression and track changes over the pseudo time.

### 2.6 RNA velocity analysis

We used the Python script velocyto to recalculate the spliced and unspliced reads for the RNA velocity analysis. py (https://github.com/velocyto-team/velocyto.py) is located in the output folder of Cell Ranger. We used velocyto. R (v0.6) from the R package to calculate the RNA velocity values for each gene in each cell, embedded the RNA velocity vector in a low-dimensional space, and mapped the velocity field on the t-SNE embedding obtained with Seurat.

### 2.7 Cell-cell communication analysis

We use CellPhoneDB (v2.0) for scRNA-seq data to identify biologically relevant ligand-receptor interactions. To identify biologically relevant ligand-receptor interactions in single-cell transcriptome data (scRNA-seq), we used CellPhoneDB (v2.0). We defined a ligand or receptor as being expressed in a specific cell type if 10% of the cells of that type had non-zero reads for the ligand/receptor encoding gene. Statistical significance was then assessed by randomly shuffling the cluster markers for all the cells and repeating the aforementioned steps, such that there was an asymmetric zero distribution for each LR in each paired comparison in both cell types. After running 1,000 permutations, we calculated the p-values using the normal curves generated from the interaction scores of the permuted LR pairs. By linking any two cell types where the ligand is expressed in the former cell type, and the receptor is expressed in the latter, we defined the intercellular communication network. We used the R package lgraph and Circlize to visualize the intercellular communication network.

### 2.8 Flow cytometry

On post-SCI day 1, we dissected the mice and removed the spinal cord after injury. After making a single-cell suspension of the injured spinal cord, bovine serum protein (5% BSA) was added to block non-specific antibody binding, and the cells were then stained for FACS according to standard procedures. We used flow cytometry (FACSCanto) and FACS Diva software to analyze the cells, and the results were processed using FlowJo software(v10.0.7).

### 2.9 Nissl and hematoxylin and eosin (HE) staining

Briefly, paraffin sections were left to dry at room temperature, and the sections were dewaxed in xylene. The sections were then removed from xylene and soaked in 100%, 95%, and 70% ethanol for 5 min. The sections were stained, sealed, and observed according to the kit instructions (Solarbio, Cat#G1434, CN). Staining was performed using the HE staining kit (Solarbio, Cat#G1120, CN), and dewaxing was performed as aforementioned.

### 2.10 Basso-Mouse Scale

The Basso-Mouse Scale was used to assess the functional recovery of mice on days 1, 3, 7, 14, and 28 after SCI. Scores ranged from 0 to 9 (9 being completely normal, and 0 being completely paralyzed). The mice were placed in an open area and allowed to move freely for 5 min. The mice were scored by observing their range of motion and limb coordination. Three examiners, who did not know the grouping of the mice, observed and scored the mice. The experiment was repeated thrice.

### 2.11 Data and code availability

The supplementary material provides additional data to support the conclusions of this paper. All the raw data and processed scRNA-seq files were stored in the GEO database under the login code GEO:#[pending]

### 2.12 Statistical Analysis

RNA sequencing data were analysed in R as described in the Methods section. For the remainder, we used Graphpad Prism 8(v 8.0.2) for statistical analysis.

## 3. Results

### 3.1 Effect of zinc therapy on mice motor function

Our previous study showed that zinc treatment could affect functional recovery in mice with spinal cord injury[5,16]. Here, we used gait scores, BMS scores, HE staining, and Nissler staining to verify the effect of zinc again. The gait test showed changes in hind limb strength after seven days of zinc treatment (Fig 1A), and the BMS score showed a significant functional recovery in the zinc-treated mice compared to the SCI group from day 14 onwards (Fig 1B). Histological analysis by Nissler staining and HE staining indicated that the spinal cord injury after zinc treatment had better integrity and neuronal activity (Fig 1C, D). These results suggest that zinc treatment can promote functional recovery in mice with spinal cord injury.

**Figure 1.**
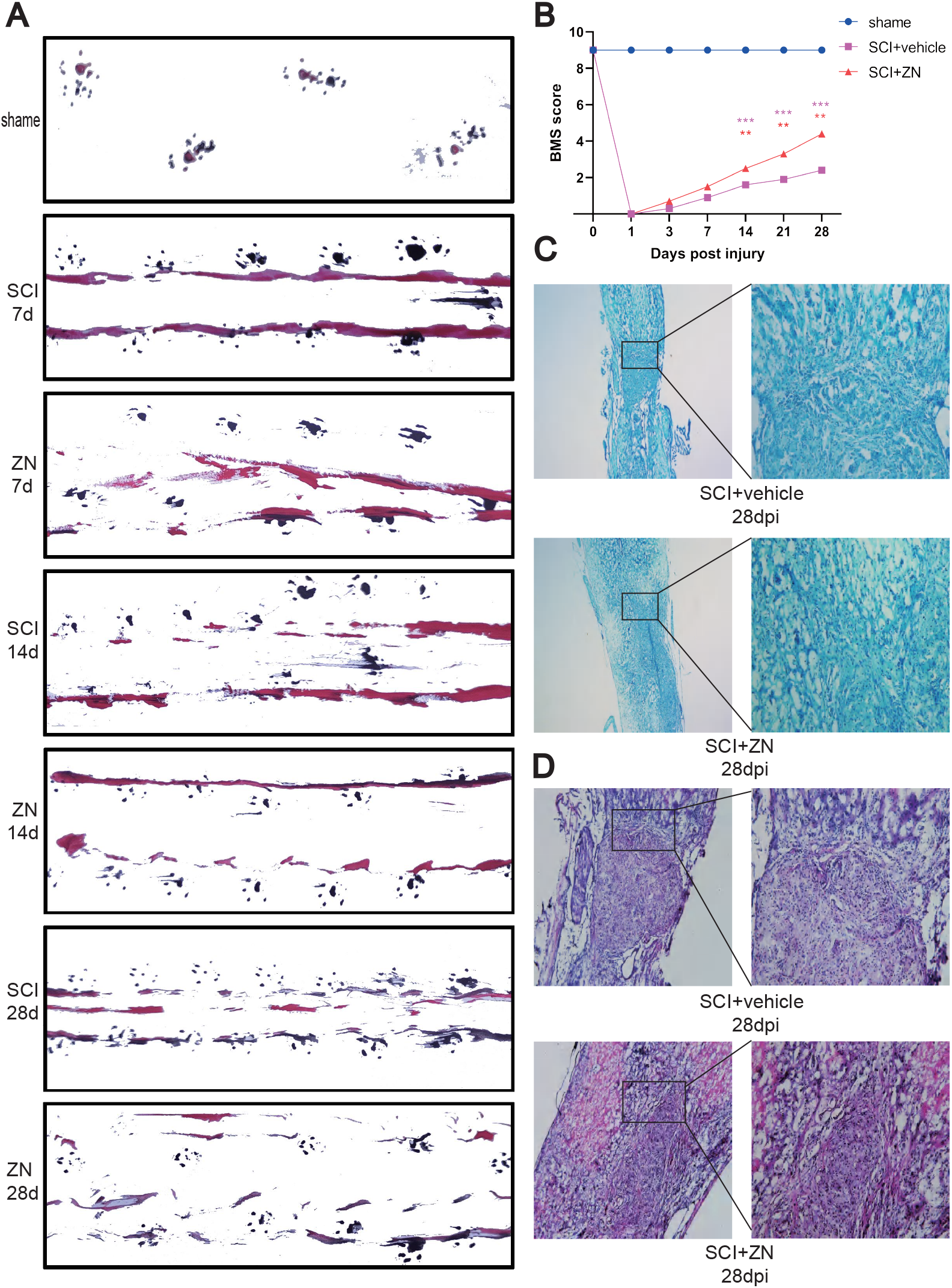
Zinc therapy promotes functional recovery after spinal cord injury (SCI) in mice. A: Gait analysis of mice at different time periods after SCI (7 dpi, 14 dpi, 28 dpi, N = 6 mice/group) B: Basso-Mouse Scale scores of mice (N = 10mice/group, all the data are expressed as means ± SD, *p<0.05,**p<0.01, ***p<0.001,****p<0.0001.) after SCI C: Hematoxylin and eosin staining of the spinal cord tissue in mice (N = 6 mice/group) 28 days after SCI D: Nissl staining of the spinal cord tissue in mice (N = 6 mice/group) 28 days after SCI. Representative mice from each group of conditions were selected for display.

### 3.2 scRNA-seq results of the cell populations of the spinal cord after SCI and zinc therapy

To generate a comprehensive single-cell atlas of the spinal cord, we isolated 20 spinal cord samples (10 each in the simple injury and zinc therapy groups) approved for research purposes. The tissues were dissociated into single cells and subjected to 10× Genomics platforms for scRNA-seq (Fig. 2A). Following strict quality control (Fig. S1, Fig. S2), a total of 30,078 cells were selected and divided into two groups: the SCI group, including an average of 1,990 genes and 14,388 transcripts per cell; and the zinc group, including an average of 1,252 genes and 15,690 transcripts per cell (Fig. S1A, B). On unsupervised clustering, we divided all cells into 18 clusters by the student t-distributed t-SNE (Fig. 2B). According to the heatmap of the top 10 genes (Fig. 2C), we identified 11 cell types (Fig. 2D) and found the expression identification of each cell type (Fig. S3A–L). These include monocytes with the typical marker genes CCL17(clusters 12 and 16); T cells with the specific marker genes CD3D, CD3E, CD3G, CD4 and CD8A (cluster 11); neurons with the distinct marker genes GAP43 and MAP2 (cluster 10); endothelial cells with the typical marker gene EGF17 (cluster 8); oligodendrocytes with the specific marker genes Olig1 and Olig2 (clusters 6 and 7); and macrophages with the distinct marker genes F4/80, CD68 and S100a4 (clusters 2 and 9); microglia with specific marker genes TMEM119, CX3CR1 (clusters 3, 4); neutrophils with specific marker genes CXCR2, LTB (clusters 1, 5); B cells with specific marker genes CD19, CD79A and CD79B (cluster 13); astrocytes with specific marker genes GFAP and AQP4 (cluster 15) and fibroblasts with typical marker genes DPEP, MME (clusters 14, 17). The distribution of cell types in each sample did not differ significantly, suggesting that cell aggregation was caused by physiological differences rather than variations arising from the technical batch or individual differences.

**Figure 2.**
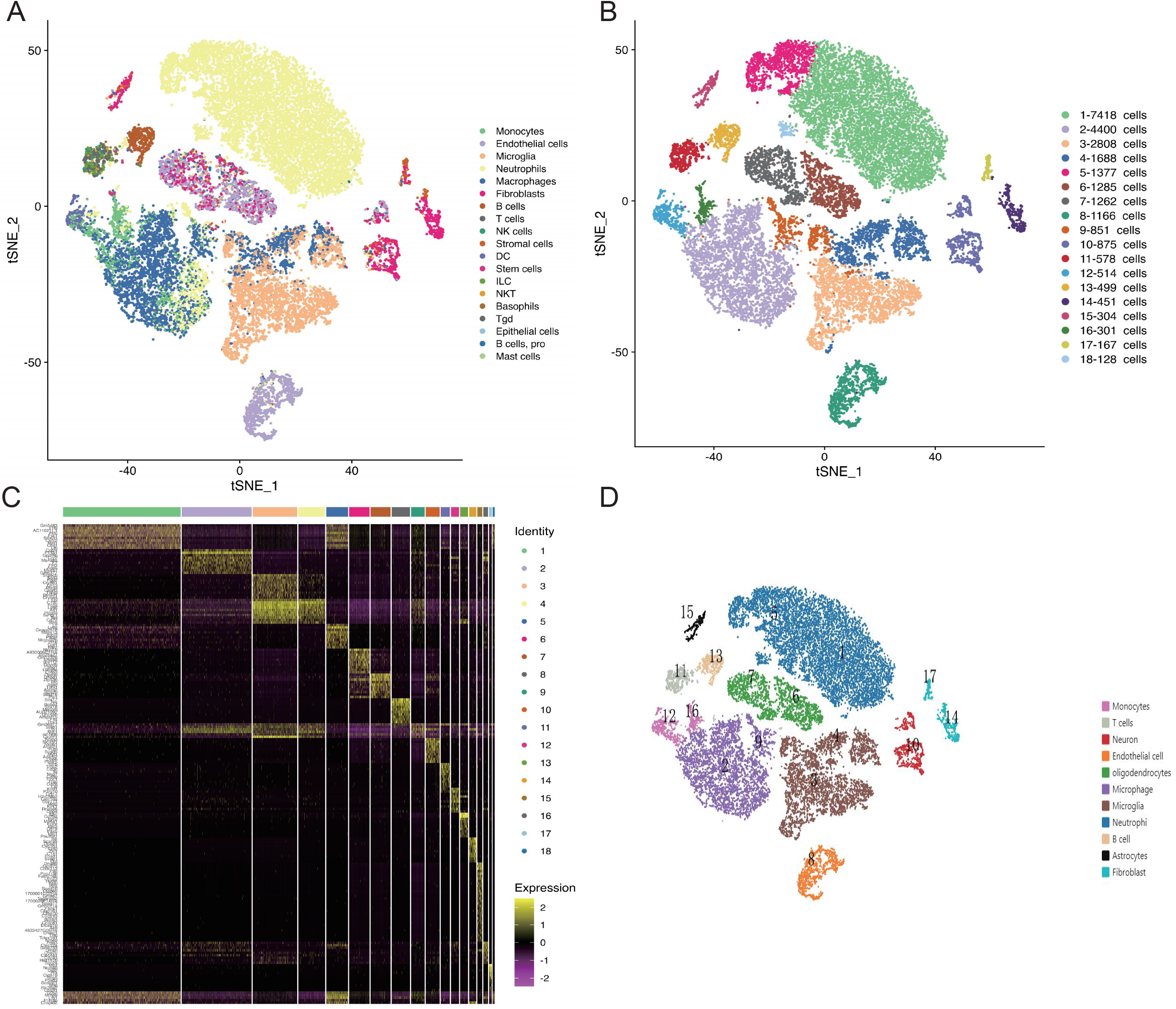
Single-cell RNA sequencing identifies different cell populations in the spinal cord tissue following spinal cord injury and zinc therapy. A: Preliminary results of single-cell RNA sequencing on mice(N=10mice/group); B: Student t-distributed stochastic neighbor embedding for visualization of cells in isolated mouse spinal cord tissue(adjusted p-value < 0.05); C: Heatmap demonstration of the top 10 genes in the mouse spinal cord tissue(adjusted p-value < 0.05); D: Student t-distributed stochastic neighbor embedding display of the 11 cell types identified(adjusted p-value<0.05)

### 3.3 Heterogeneous changes in the injured spinal cord caused by zinc

Based on the credible single-cell atlas of the spinal cord, we continued to explore the heterogeneous effects of zinc ion therapy on the spinal cord. We analyzed the changes in cell clusters in the simple injury and zinc therapy groups by using t-SNE (Fig. 3A). The difference in the cell clusters between the SCI and zinc groups was obtained utilizing data visualization (Fig. 3B–D). The results showed significant quantitative differences in neutrophils, macrophages, and microglia; populations after zinc therapy. In terms of the number of cells, neutrophils showed chemotactic aggregation (62.31% zinc vs. 37.69% SCI), with a small increase in the microglia (54.25% zinc vs. 45.75% SCI) and reduction in macrophages (46.26% zinc vs. 53.74% SCI). Meanwhile, by analyzing the differential genes between the SCI and zinc groups, we identified 644 genes with differential properties. Among them, the expression of 487 genes was up-regulated, and 157 genes were down-regulated in the zinc group compared to the SCI group. Also, after comparing the zinc and SCI groups, the results of GO and KEGG analysis of TOP30 differential genes in the zinc group indicated that zinc treatment is involved in biological inflammatory processes (Figure 3E, F). This appears to be associated with post-infection signal transduction pathway activation, such as lysosomal and IL-17 activation. These results suggest that a daily therapeutic dose of zinc therapy affects not only the number of chemotactic inflammatory and innate immune cells in the injured spinal cord but also the immune function in the spinal cord microenvironment.

**Figure 3.**
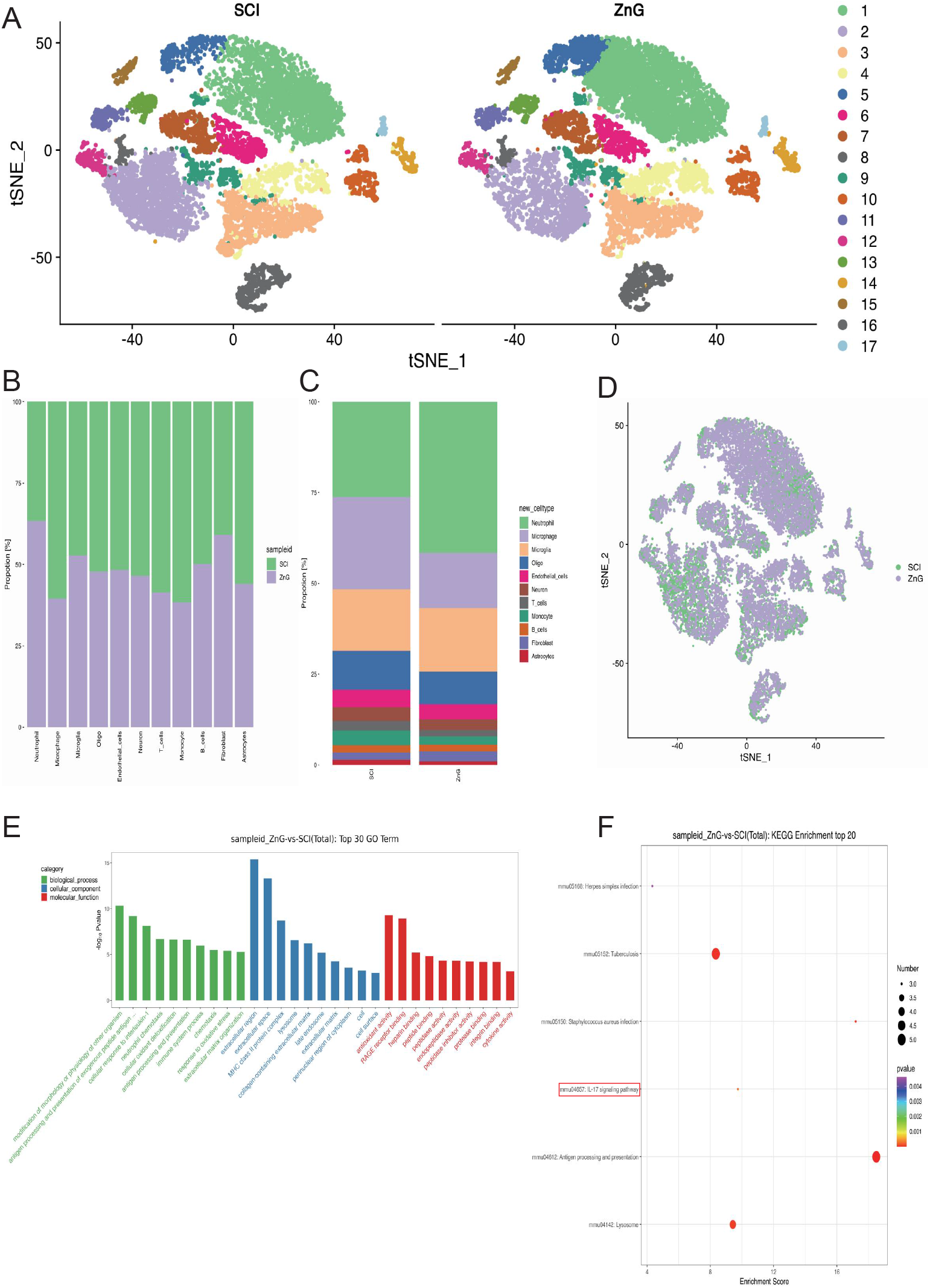
Zinc therapy leads to heterogeneous changes in spinal cord tissue after spinal cord injury. A: Student t-distribution stochastic neighbor embedding showing the difference in cell clusters between the treatment-only and zinc therapy groups(adjusted p-value<0.05); B–D: Different data visualization methods to examine cell cluster differences between the spinal cord injury and zinc groups(adjusted p-value<0.05); E: Gene oncology analysis suggesting that zinc therapy is involved in the inflammatory process; F: KEGG analysis suggesting that zinc therapy is involved in the inflammatory process

### 3.4 Microglia classification based on cell type-specific marker genes

Based on the already obtained plausible differences in cell clusters between the SCI and zinc treatment groups, we first focused on the main immune cells involved in the inflammatory response in the injured spinal cord, i.e. microglia. To further elucidate the effect of zinc therapy on inflammatory pathological processes in the spinal cord microenvironment, re-clustered and performed pseudo-temporal trajectory analysis. Again, further analysis of shared nearest-neighbour clustering of microglia using the single-cell R Toolkit Seurat yielded seven juxtaposed clusters (M1, 28.67%; M2, 16.74%; M3, 16.27%; M4, 12.17%; M5, 12.13%; M6, 10.49%; M7, 2.1%; Fig. 4A). We further identified these subclusters based on the expression levels of cell marker genes and obtained marker gene heatmaps for each subpopulation (Fig. 4B). Based on the gene heatmap for each cluster, we clustered the juxtaposed subgroups into five subgroups (Fig. 4C): Clusters 1 and 5 expressed relatively high levels of the proliferation-related gene Mki67, which we defined as the Mki67+ microglia, and Clusters 2 and 3 expressed relatively high levels of Ppia and ribosome-related genes, which we defined as the Ppia Microglia. Cluster 4 was defined as the Vegfa+ microglia. Cluster 6 expressed high levels of lfit2 and Bbc3, which we defined as the lfit2+ microglia. Cluster7 expressed high-level genes Ms4a7, Cd163, and Lye1, which we defined as the Ms4a7+ microglia (Fig. 4D).

**Figure 4.**
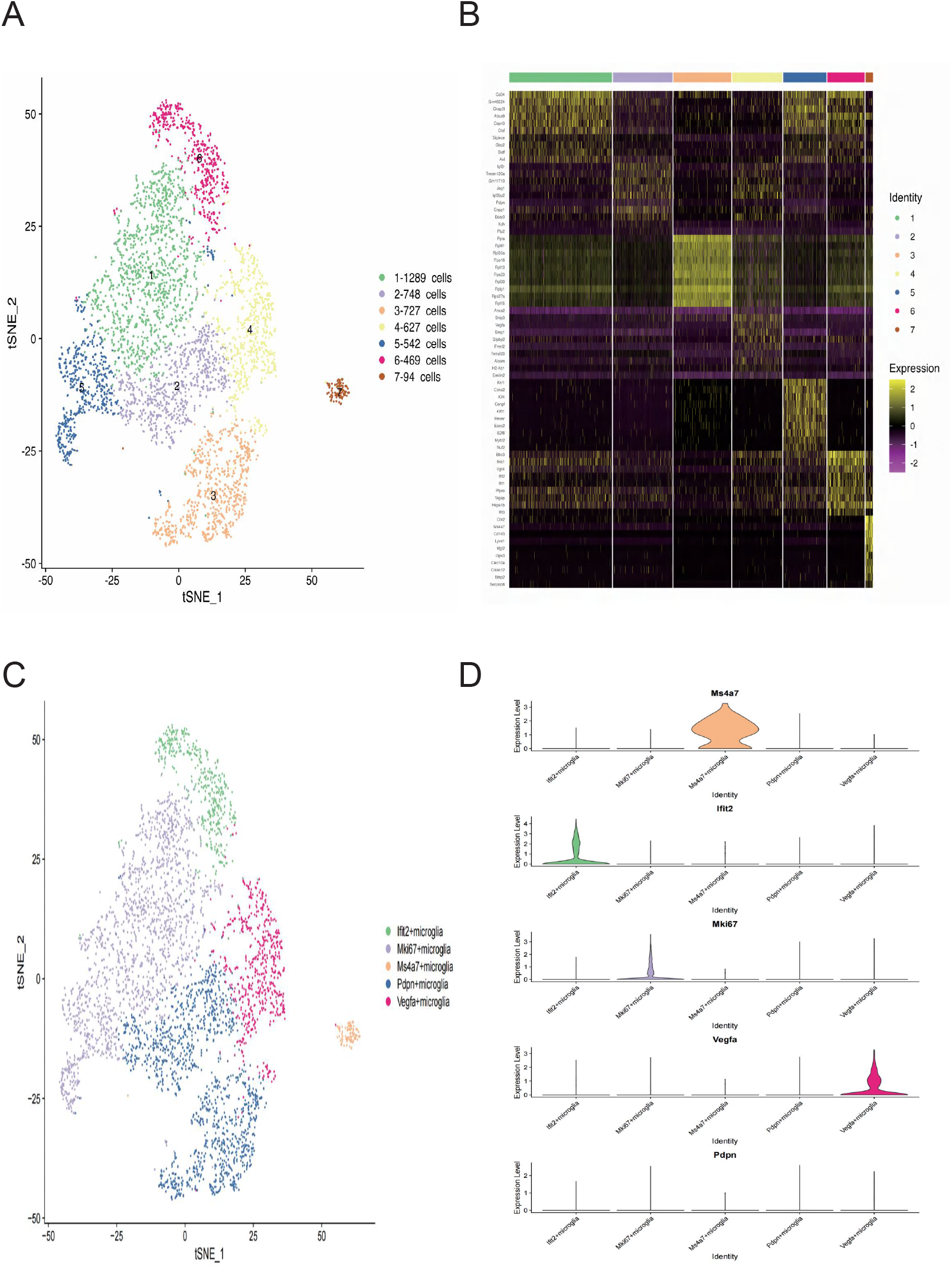
Microglial cell populations based on different marker genes. A: Nearest-neighbor class analysis of the microglia; B: Heatmap showing marker genes for various subpopulations of the microglia(adjusted p-value<0.05); C-D: Grouping of microglia subpopulations.

### 3.5 Reconstructing the developmental trajectory of microglia in spinal cord tissue after zinc treatment

The pseudotime trajectory analysis of the progression of successive cell states in the microglia showed ordered cells expressing different levels of marker genes in a single trajectory (Fig. 5A). The order of pseudo-time cells arranged most glial cells into a major trajectory with two bifurcations and nine stages (Fig. 5A, S4). The proliferative subcluster Mki67+ microglia was located towards the origin of the trajectory, which partly served as a validation for the constructed trajectory. The subcluster Ppia Microglia, which expresses more ribose genes, serves as the end point of the trajectory (Fig. 5B). Early (Mki67+ microglia) and late (ribosome+ microglia) microglia were located at the end of the trajectory bifurcation, while the angiogenic phenotype (Vegfa+ microglia) and Ms4a7+ microglia were located at the other branch end of the trajectory (Fig. 5A, B). We examined the pseudo-temporal dynamics of genes that underwent significant changes in these five subpopulations of cells and divided their pseudo-temporal expression patterns into three modules (Fig. 5C).To reshape cellular differentiation in all microglia, we performed the RNA velocity analysis to deconstruct changes in RNA dynamics and direction of state transitions and found that Mki67 microglia exhibited lower RNA velocities (short or no arrows) in the quiescent state. The RNA in the Vegfa and ribosomal microglia was fast and consistent with the initiation of cell differentiation (Fig. 5D). These results corroborate the pseudo-time analysis results.

**Figure 5:**
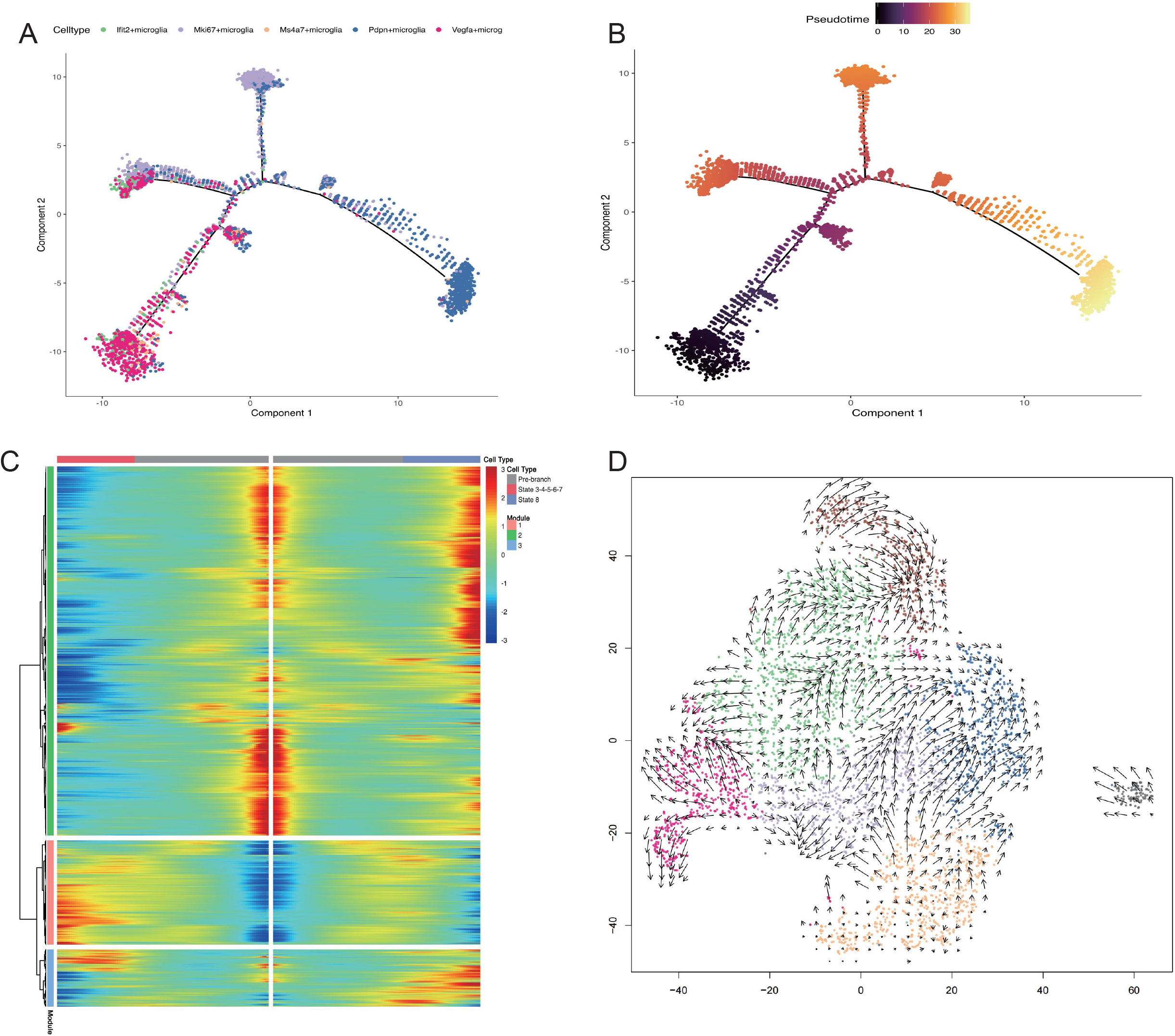
Reconstructing the developmental trajectory and the pseudotime analysis of microglia. A: Monocle2-generated pseudotemporal trajectory of 7 microglia subtypes imported from Seurat data, coloured by cell-name designation; B: Pseudotime was colored in a gradient from dark to light red, and the start of pseudotime was indicated; C: Heat map for clustering the top 50 genes that vary as a function of pseudotime. The 50 genes were divided into three clusters (cluster 1, cluster 2, and cluster 3), representing the genes at the beginning stage, the transitory stage, and the end stage of developmental trajectory, respectively; D: Pseudotime analysis of microglia by using RNA velocity reconstructed the developmental lineages.

### 3.6 Protein interaction networks reveal neuronal changes in the immune microenvironment following zinc treatment

To further understand the specific effects of the microglia in the immune microenvironment after zinc therapy, we constructed a cell-cell communication map using intercellular ligands and receptors (Fig. 6A). By counting the number of ligand-receptor binding sites (Fig. 6B), we visualized the results obtained using a matrix heatmap (Fig. 6C) and found that the microglia were closely associated with the macrophages and monocytes, suggesting that the glial cells were more active under zinc therapy (Fig. 6C). Next, we performed a specific analysis of the ligand-receptor binding sites of the microglia (Fig. 6D). Zinc therapy altered the communication between the microglia and other cells. In most cases, zinc had an antagonistic effect. Tgf-beta1, Csf1, and Sema4d signalling pathways were significantly decreased, while IL1 and Tnf-a signalling pathways were significantly increased in the interaction between the macrophages and the microglia (Fig. 6D). These results suggest that zinc therapy enhanced the viability of the microglia and significantly enhanced their interaction with immune cells, such as macrophages and monocytes.

**Figure 6.**
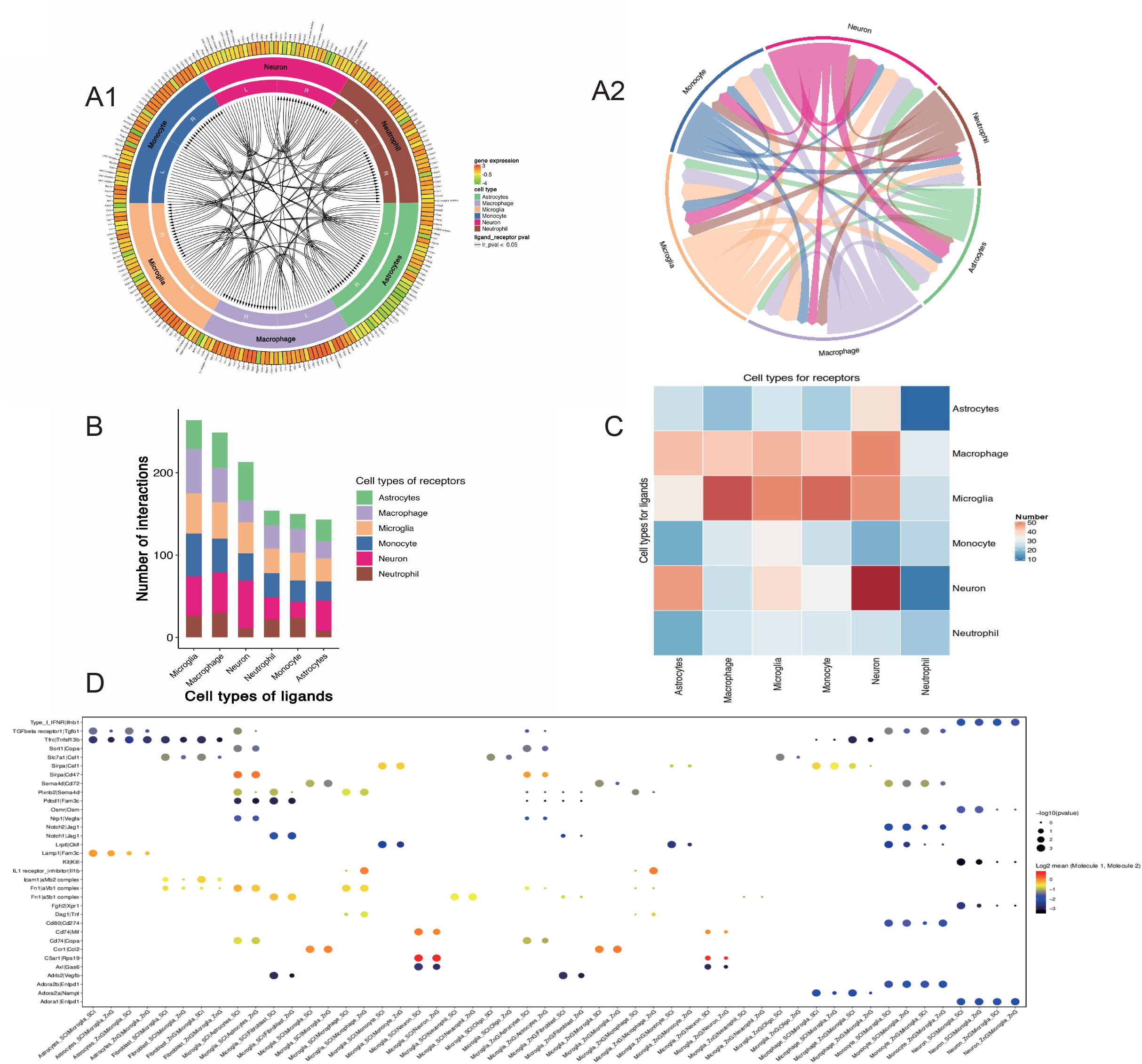
Protein interaction networks reveal neuronal changes in the immune microenvironment owing to zinc therapy. A: Communication map among cells(adjusted p-value<0.05); B: Calculation of receptors and ligands between cells(adjusted p-value<0.05); C: Data visualization of receptor and ligand calculations between cells(adjusted p-value<0.05); D: Specific analysis of microglia ligand receptor binding sites(adjusted p-value<0.05)

### 3.7 Identification of a novel VEGFA+ microglia cell type

Using flow cytometry, we extracted microglial populations and validated gene expression across subpopulations (Fig. 7A-H). The results of the pseudo-chronological analysis of the microglia revealed the effect of immune activation produced by zinc on angiogenesis. To observe the effect of zinc therapy on the reconstitution of blood flow to the injury site after SCI, we performed immunofluorescence staining for CD31 on the spinal cord from mice 28 days after SCI. The results showed a significant increase in CD31-positive cells in the SCI + zinc group compared to the SCI group (Fig. S5), suggesting that zinc therapy significantly increased revascularization and regeneration at the injury site compared to the SCI group. The results indicate that zinc therapy can promote vascular regeneration at the injury site by activating the microglia.

**Figure 7.**
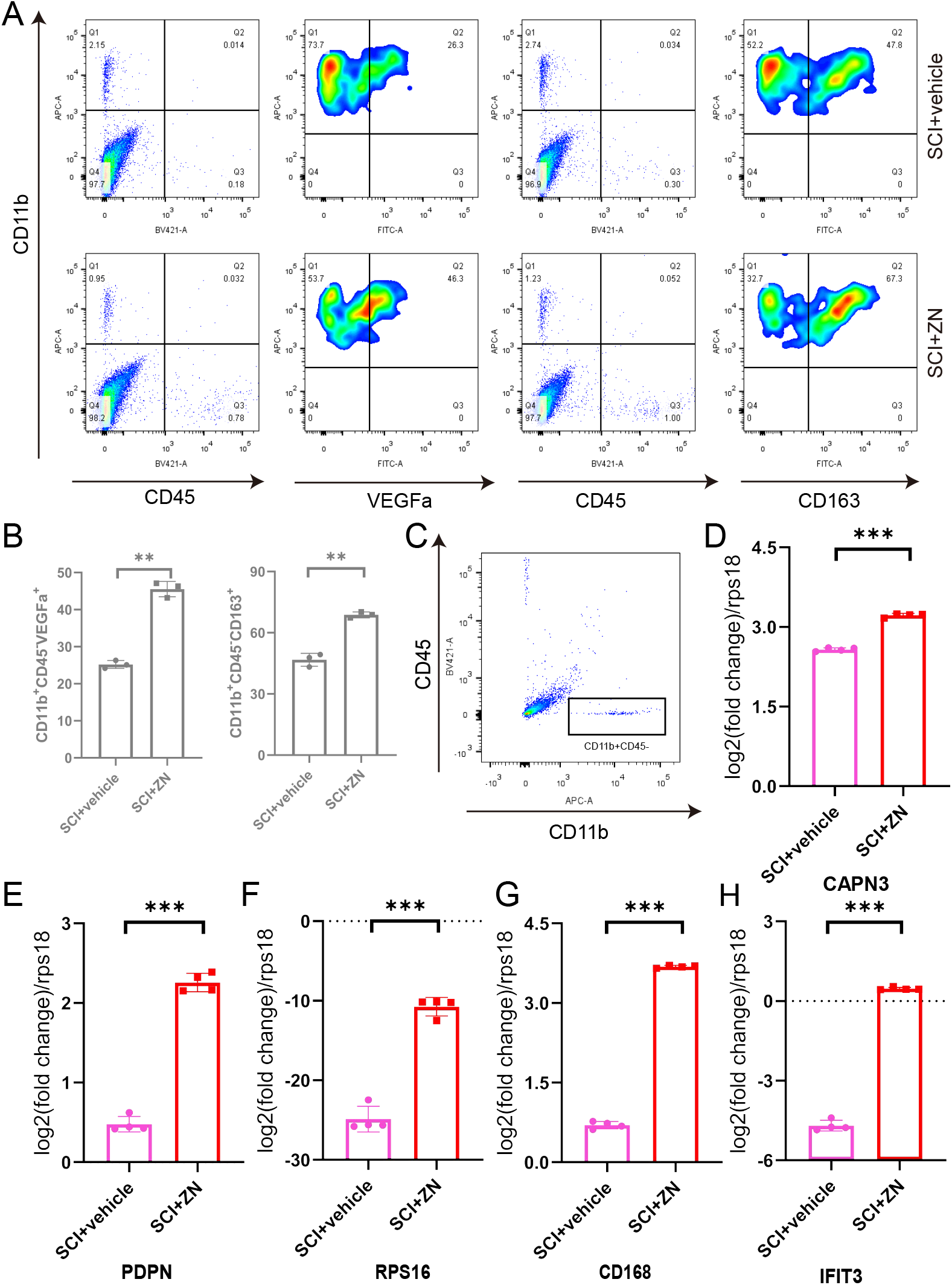
Identification of microglial cell subpopulations. A, B: VEGFA_+_ and CD163_+_ microglia subpopulations in the SCI and SCI+ZnG groups were analyzed and identified using flow cytometry(N=3 mice/group). C: Application of flow cytometric sorting techniques to identify microglia. D-H: Validation of gene expression in the microglia using flow cytometry(N=4 mice/group). Representative mice from each group of conditions were selected for display.*p<0.05,**p<0.01, ***p<0.001,****p<0.0001;

## 4 Discussion

SCI is a serious central nervous system injury disorder[28]. Severe SCI can cause quadriplegia or even high-level paraplegia, severely affecting a patient’s quality of life[29]. SCI include primary and secondary injuries, with primary injuries often occurring within a few hours and secondary spinal cord injuries often lasting for weeks or even months[30]. The pathophysiology of secondary SCI places particular emphasis on endothelial cell dysfunction and altered vascular permeability due to ischaemia-reperfusion, which can lead to a cascade of inflammatory responses, resulting in the activation of astrocytes and microglia and infiltration of neutrophils and macrophages in the spinal cord tissue[31–34]. These recruited inflammatory cells release an abundance of inflammatory factors and chemokines, further exacerbating the inflammatory response, resulting in neuronal death and loss of neurological function in the patient[35–40]. The therapeutic approach to SCI emphasises individualised and comprehensive treatment, with the key being specific interventions at different stages of the inflammatory process[41]. Our study showed that improvement in motor function occurred in mice with SCI after 28 days of treatment with zinc, suggesting that zinc could be used as a potential treatment for SCI.

In previous studies, immune cells are involved in functional recovery after SCI, and microglia also appear to have functions other than immune functions, but the specific mechanisms involved have not been explored[23,42,43]. Therefore, we focused on microglia as intrinsic immune cells in the CNS. By performing a shared-neighbourhood clustering analysis of zinc-treated microglia via the single-cell R toolkit Seurat, we created seven subclusters of microglia. In annotating the subclusters of microglia with highly expressed genes, we found that microglia appear to be immunologically heterogeneous. We then grouped the seven microglia subclusters into five cell subtypes based on specific highly expressed genes. In parallel, we performed microglia sorting by flow cytometry and verified the presence of subpopulations.

In contrast to previous knowledge, microglia are no longer classified as purely anti-inflammatory phenotype M2 and purely pro-inflammatory phenotype M1[33,44-48]. Our proposed time-series analysis shows that Ki67+ microglia have proliferative functions and may differentiate into other functional microglia, such as Vegfa+ microglia. This finding suggests that similar to previous reports, microglia may be involved in microangiogenesis at the site of early spinal cord injury and may also play a neuroprotective role in the early stages of spinal cord injury[49–52], which may be enhanced by zinc treatment. Furthermore, in an in vitro model, we observed high expression of CD31 in tissues at the site of spinal cord injury on day 28 after zinc treatment, suggesting that zinc could influence angiogenesis by affecting the inflammatory response in the central nervous system. Using protein-protein interaction networks, we analysed the association of ligands and receptors with microglia and found that zinc treatment could affect the ‘crosstalk communication’ of various cells in the SCI tissue microenvironment. Firstly, neuronal interactions are very close, which may be related to the high expression of zinc finger proteins in neuronal cells, allowing for better zinc uptake. Secondly, microglia are closely associated with peripherally derived chemotactic neutrophils, macrophages and monocytes, while they are associated with astrocytes and less so with oligodendrocytes in the nervous system. This phenomenon was not explored in this study, but we will continue to do so in subsequent studies.

In summary, we describe microglia dynamics and transcriptome profiles in zinc-treated SCI by single-cell sequencing techniques. We show that short-term zinc therapy can the immune microenvironment after SCI, which may be associated with activation of the IL-17 signalling pathway. Meanwhile, long-term zinc therapy promotes postoperative vascular regeneration in SCI mice, which may be associated with the differentiation of functional microglia. Thus, our study could serve as a molecular basis for zinc therapy in the treatment of SCI and may help rationalize the clinical use of zinc in the treatment of SCI.

## ACKNOWLEDGMENTS

This work was supported by the National Natural Science Foundation of China (NSFC; Grant NO.81671907, NO.81871556), LiaoNing Revitalization Talents Program (No.XLYC1902108), and the Scientific Research Project of the Educational Department of Liaoning Province (No.JYTQN201917).

## AUTHOR CONTRIBUTIONS

Z.S and J.L have made equal contributions to this work. Z.S, J.L, and L.M completed the entire conception and design of the experiment. Z.S, J.L, M.G, and H.H were engaged in behavioural scoring and sample preparation. Z.S, J.L, M.G, and H.H were involved in HE and Nissl staining. Z.S, J.L, and M.T concluded the statistical analysis and manuscript preparation. X.M and H.T Professors completed the final review and presented the manuscript. All authors contributed significant intellectual content to the manuscript and gave their consent for publication.

## Figure Legends

**Figure S1:**
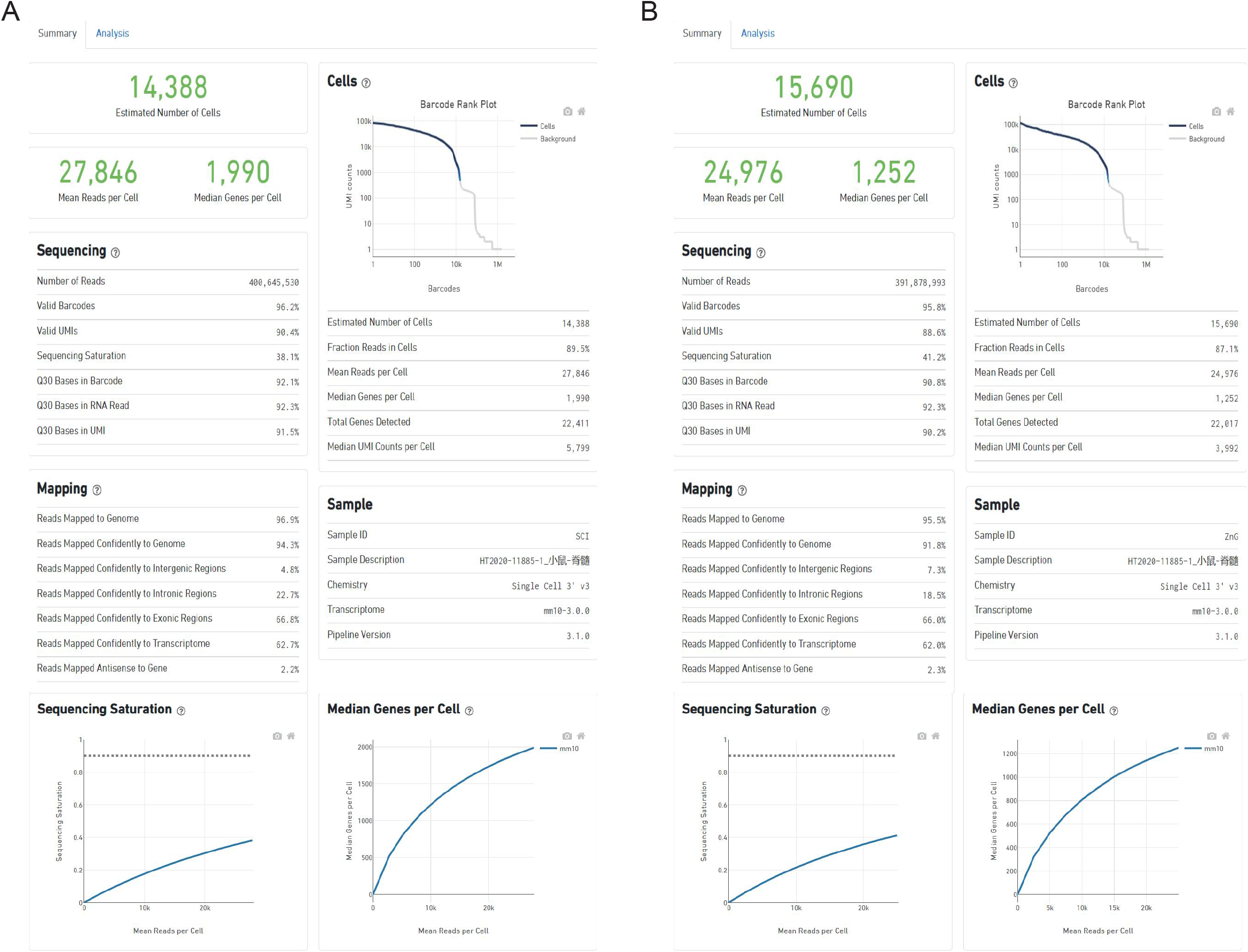
summary of single-cell sequencing results. A: The SCI group; B: The SCI+ZnG group.

**Figure S2:**
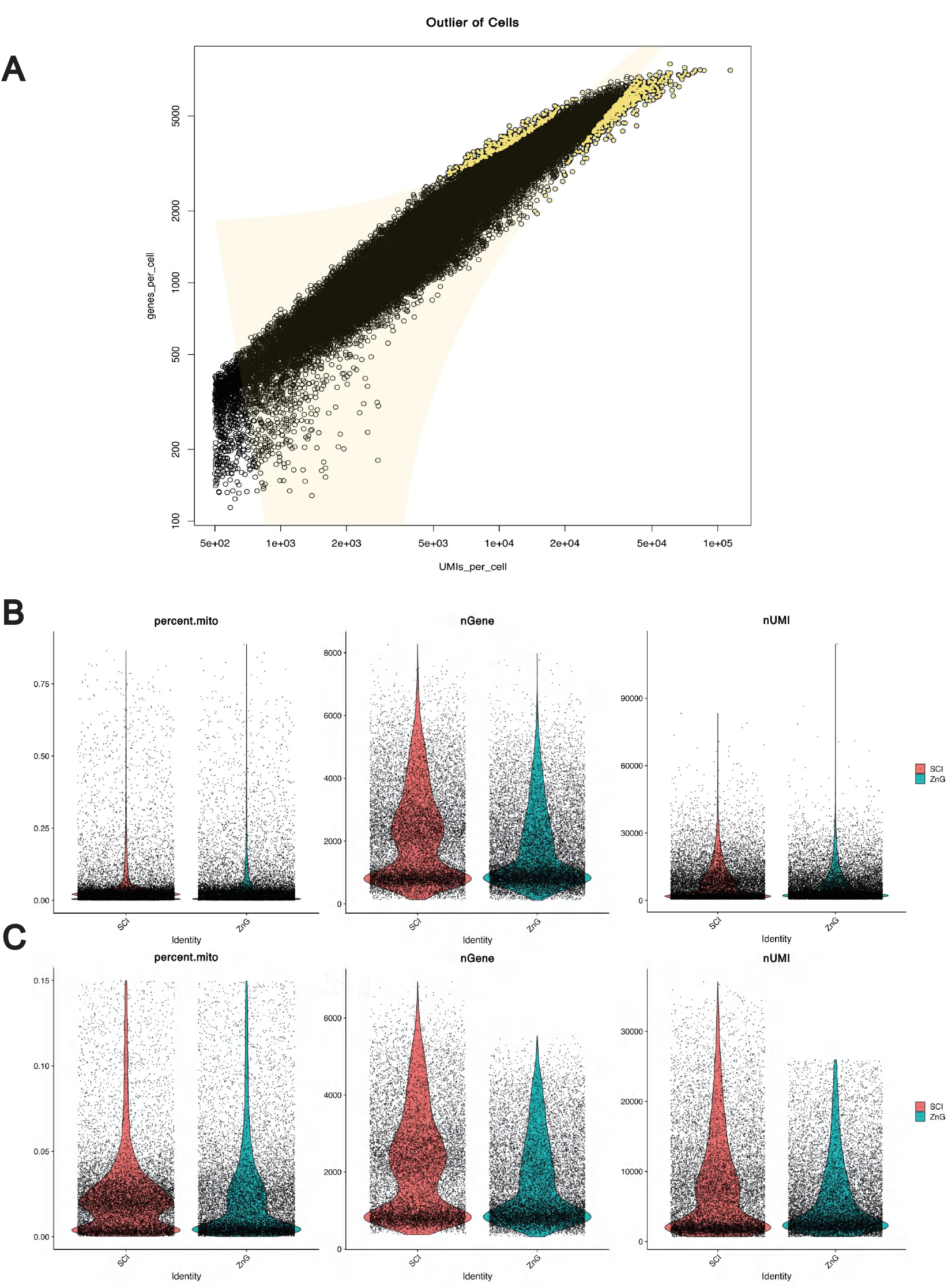
Quality control of single-cell sequencing results. A: Filter out the normal individual live cells; B: Distribution of mitochondrial gene transcripts, number of genes and number of UMIs in individual cells in single cells before quality control; C: Distribution of mitochondrial gene transcripts, number of genes and number of UMIs in individual cells in single cells after quality control.

**Figure S3:**
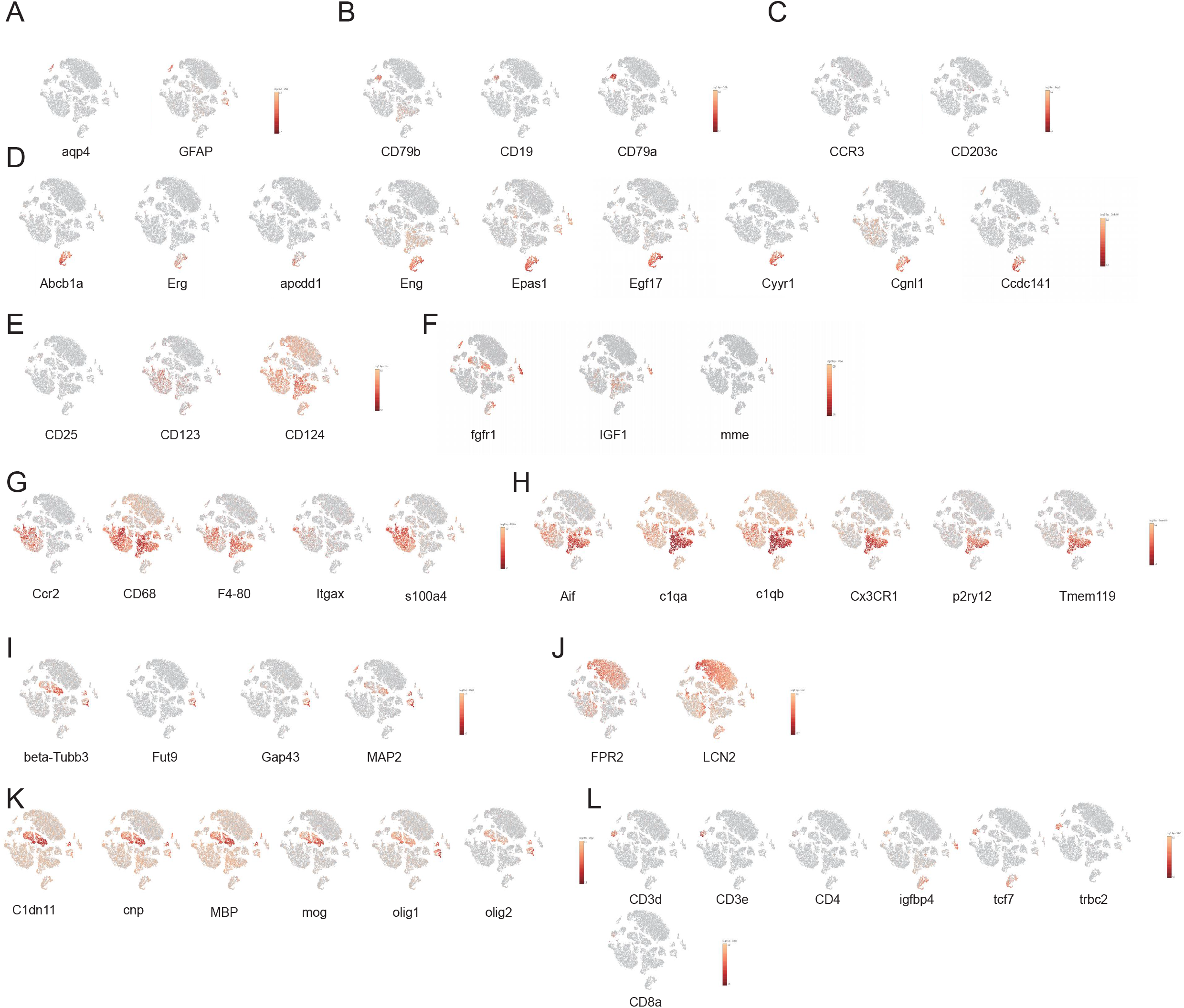
Expression markers for each cell type in sequencing results. A: Astrocyte B: B cell C: Monocytes D: Endothelial cell E: Fibroblast F: Macrophage G: Microglia H: Neuron I: Neutrophil J: Oligodendrocytes K: T cell Microglia mimetic time-series analysis revealing differentiation trajectories.

**Figure S4:**
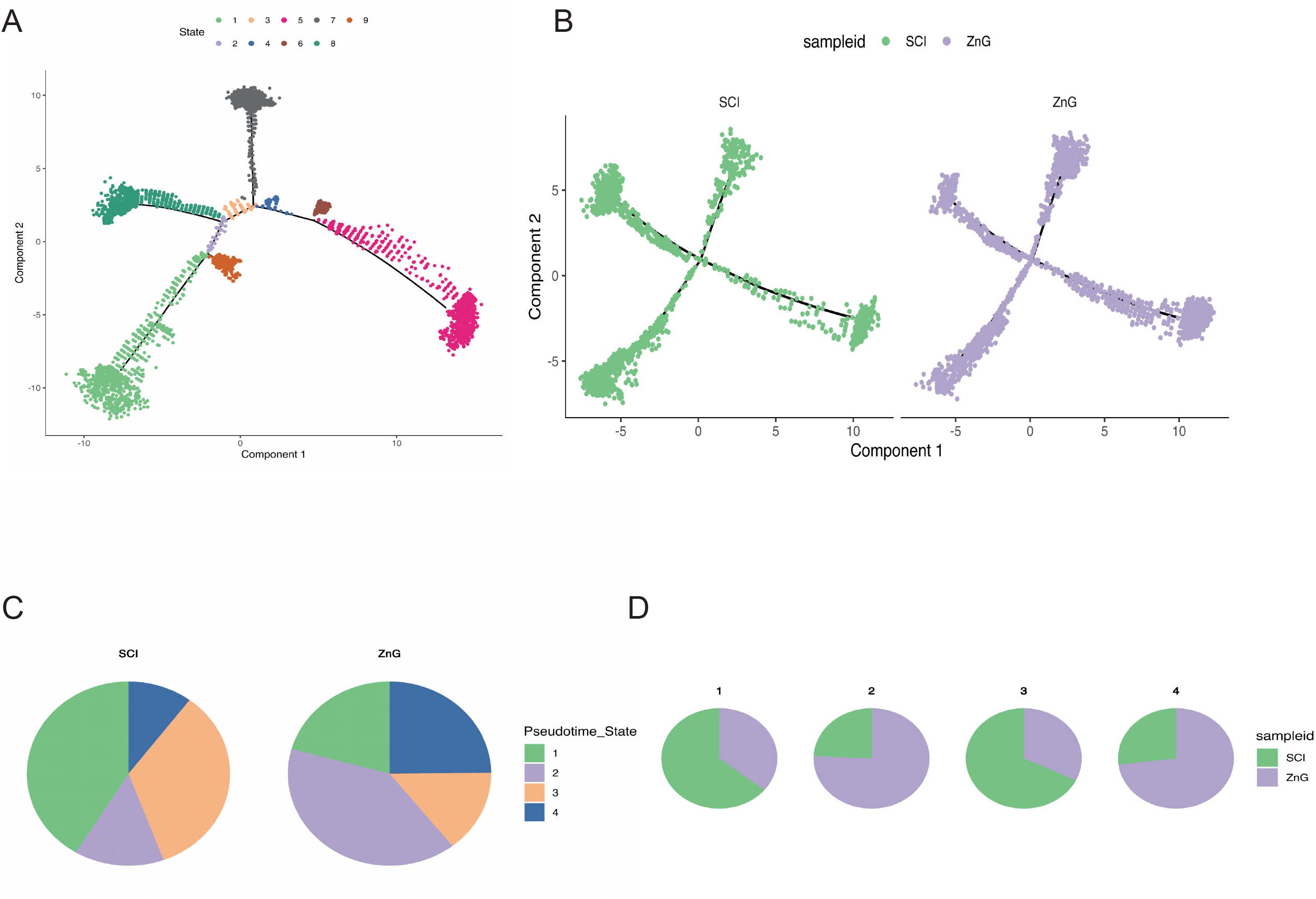
A: Distribution of microglia of different phenotypes on the differentiation trajectory; B: Comparison of the differentiation trajectories of microglia in the SCI and SCI+ZnG groups; C: Comparison between the number of cells in each state of the microglia differentiation trajectory in the SCI and SCI+ZnG groups.; D: Comparison between the number of cells in each state of the microglia differentiation trajectory in the SCI and SCI+ZnG groups.

**Figure S5:**
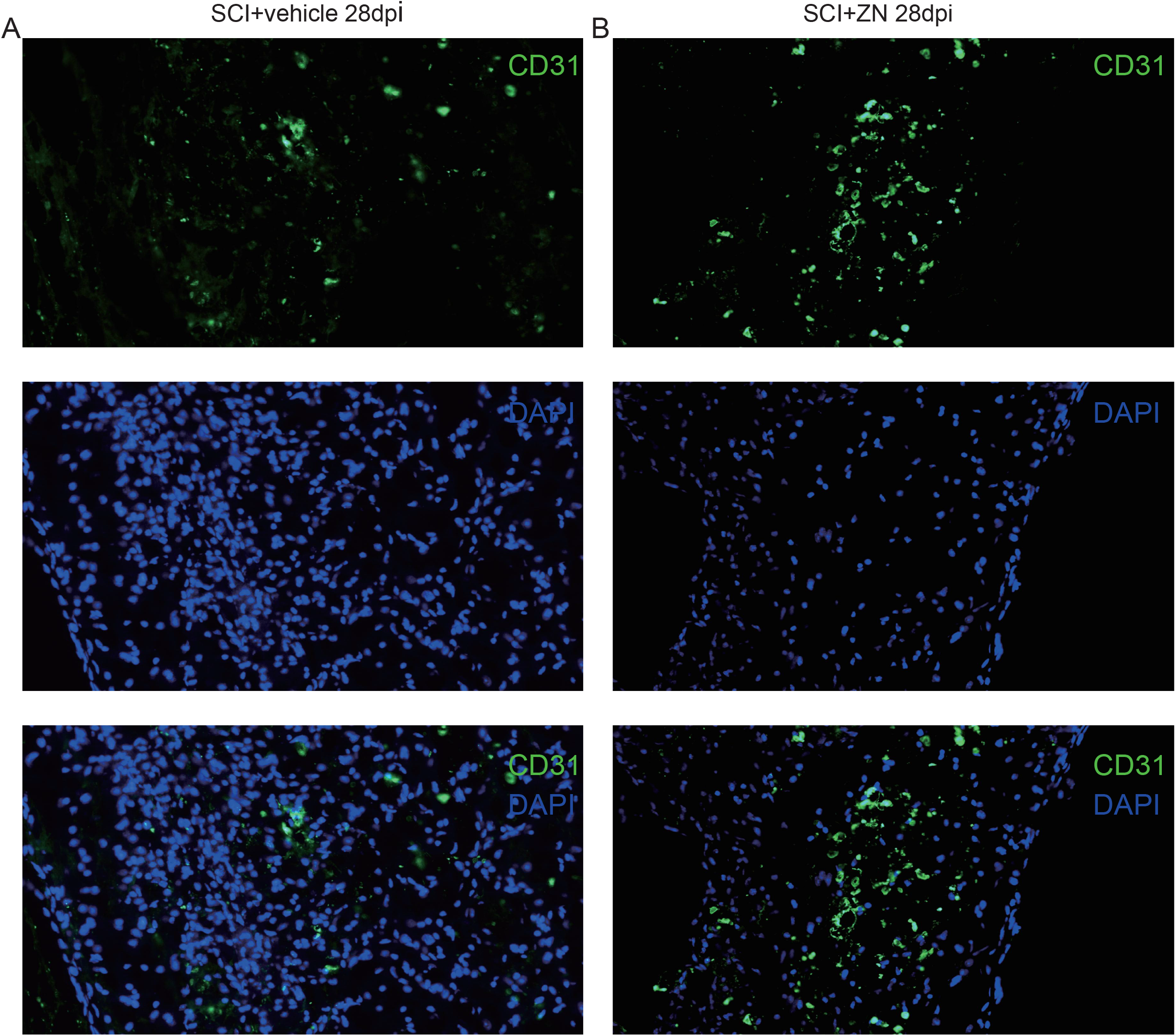
A: Immunofluorescence staining of the spinal cord tissue from the SCI group (28 dpi). The green colour shows CD31 expression B: Immunofluorescence staining of the spinal cord tissue from the SCI + zinc group(28 dpi). The green colour shows CD31 expression A representative mice from each group of conditions were selected for display(N=3mice/group).Scale bar = 100 μm.

## REFERENCES

1. Alizadeh A, Dyck SM, Karimi-Abdolrezaee S. Traumatic Spinal Cord Injury: An Overview of Pathophysiology, Models and Acute Injury Mechanisms. Front Neurol. 2019 Mar 22;10:282. doi: 10.3389/fneur.2019.00282.

2. Beattie MS. Inflammation and apoptosis: linked therapeutic targets in spinal cord injury. Trends Mol Med. 2004 Dec;10(12):580–3. doi: 10.1016/j.molmed.2004.10.006. PMID: 15567326.

3. Rowland JW, Hawryluk GW, Kwon B, Fehlings MG. Current status of acute spinal cord injury pathophysiology and emerging therapies: promise on the horizon. Neurosurg Focus. 2008;25(5):E2. DOI: 10.3171/FOC.2008.25.11.E2. PMID: 18980476.

4. Chan R C K. How does spinal cord injury affect marital relationship? A story from both sides of the couple[J]. Disability and Rehabilitation, 2000, 22(17): 764–775.

5. Li X, Chen S, Mao L, Li D, Xu C, Tian H, Mei X. Zinc Improves Functional Recovery by Regulating the Secretion of Granulocyte Colony Stimulating Factor From Microglia/Macrophages After Spinal Cord Injury. Front Mol Neurosci. 2019 Feb 1;12:18. doi: 10.3389/fnmol.2019.00018. PMID: 30774583; PMCID: PMC6367229.

6. Morris DR, Levenson CW. Zinc in traumatic brain injury: from neuroprotection to neurotoxicity. Curr Opin Clin Nutr Metab Care. 2013 Nov;16(6):708–11. doi: 10.1097/MCO.0b013e328364f39c. PMID: 23945221.

7. Kitamura H, Morikawa H, Kamon H, Iguchi M, Hojyo S, Fukada T, Yamashita S, Kaisho T, Akira S, Murakami M, Hirano T. Toll-like receptor-mediated regulation of zinc homeostasis influences dendritic cell function. Nat Immunol. 2006 Sep;7(9):971–7. doi: 10.1038/ni1373. Epub 2006 Aug 6. PMID: 16892068.

8. Stefanidou M, Maravelias C, Dona A, Spiliopoulou C. Zinc: a multipurpose trace element. Arch Toxicol. 2006 Jan;80(1):1–9. doi: 10.1007/s00204-005-0009-5. Epub 2005 Sep 27. PMID: 16187101.

9. Yu Z, Yu Z, Chen Z, Yang L, Ma M, Lu S, Wang C, Teng C, Nie Y. Zinc chelator TPEN induces pancreatic cancer cell death through causing oxidative stress and inhibiting cell autophagy. J Cell Physiol. 2019 Nov;234(11):20648–20661. doi: 10.1002/jcp.28670.

10. Sekler I, Moran A, Hershfinkel M, Dori A, Margulis A, Birenzweig N, Nitzan Y, Silverman WF. Distribution of the zinc transporter ZnT-1 in comparison with chelatable zinc in the mouse brain. J Comp Neurol. 2002 Jun 3;447(3):201–9. doi: 10.1002/cne.10224. PMID: 11984815.

11. Kambe T, Hashimoto A, Fujimoto S. Current understanding of ZIP and ZnT zinc transporters in human health and diseases. Cell Mol Life Sci. 2014 Sep;71(17):3281–95. doi: 10.1007/s00018-014-1617-0. Epub 2014 Apr 8. PMID: 24710731.

12. Vyas Y, Jung Y, Lee K, Garner CC, Montgomery JM. In vitro zinc supplementation alters synaptic deficits caused by autism spectrum disorder-associated Shank2 point mutations in hippocampal neurons. Mol Brain. 2021 Jun 24;14(1):95. doi: 10.1186/s13041-021-00809-3. PMID: 34167580; PMCID: PMC8223320.

13. Morris DR, Levenson CW. Zinc in traumatic brain injury: from neuroprotection to neurotoxicity. Curr Opin Clin Nutr Metab Care. 2013 Nov;16(6):708–11. doi: 10.1097/MCO.0b013e328364f39c. PMID: 23945221.

14. Wellinghausen N, Kirchner H, Rink L. The immunobiology of zinc. Immunol Today. 1997 Nov;18(11):519–21. doi: 10.1016/s0167-5699(97)01146-8. PMID: 9386346.

15. Wellinghausen N, Martin M, Rink L. Zinc inhibits interleukin-1-dependent T cell stimulation. Eur J Immunol. 1997 Oct;27(10):2529–35. doi: 10.1002/eji.1830271010. PMID: 9368606.

16. Li D, Tian H, Li X, Mao L, Zhao X, Lin J, Lin S, Xu C, Liu Y, Guo Y, Mei X. Zinc promotes functional recovery after spinal cord injury by activating Nrf2/HO-1 defense pathway and inhibiting inflammation of NLRP3 in nerve cells. Life Sci. 2020 Mar 15;245:117351. doi: 10.1016/j.lfs.2020.117351. Epub 2020 Jan 22. PMID: 31981629.

17. Macosko EZ, Basu A, Satija R, et al. Highly parallel genome-wide expression profiling of individual cells using nanoliter droplets. Cell. 2015;161(5):1202–1214.

18. Ayyaz A, Kumar S, Sangiorgi B, et al. Single-cell transcriptomes of the regenerating intestine reveal a revival stem cell. Nature. 2019;569(7754):121–125.

19. Park J, Shrestha R, Qiu C, et al. Single-cell transcriptomics of the mouse kidney reveals potential cellular targets of kidney disease. Science. 2018;360(6390):758–763.

20. Montoro DT, Haber AL, Biton M, et al. A revised airway epithelial hierarchy includes CFTR-expressing ionocytes. Nature. 2018;560(7718):319–324.

21. Milich LM, Choi JS, Ryan C, Cerqueira SR, Benavides S, Yahn SL, Tsoulfas P, Lee JK. Single-cell analysis of the cellular heterogeneity and interactions in the injured mouse spinal cord. J Exp Med. 2021 Aug 2;218(8):e20210040. doi: 10.1084/jem.20210040. Epub 2021 Jun 16. PMID: 34132743; PMCID: PMC8212781.

22. Tedeschi A, Dupraz S, Laskowski CJ, Xue J, Ulas T, Beyer M, Schultze JL, Bradke F. The Calcium Channel Subunit Alpha2delta2 Suppresses Axon Regeneration in the Adult CNS. Neuron. 2016 Oct 19;92(2):419–434. doi: 10.1016/j.neuron.2016.09.026. Epub 2016 Oct 6. PMID: 27720483.

23. Li Y, He X, Kawaguchi R, Zhang Y, Wang Q, Monavarfeshani A, Yang Z, Chen B, Shi Z, Meng H, Zhou S, Zhu J, Jacobi A, Swarup V, Popovich PG, Geschwind DH, He Z. Microglia-organized scar-free spinal cord repair in neonatal mice. Nature. 2020 Nov;587(7835):613–618. doi: 10.1038/s41586-020-2795-6. Epub 2020 Oct 7. PMID: 33029008; PMCID: PMC7704837.

24. Lu M, Quan Z, Liu B, Jiang D, Ou Y, Zhao J. Establishment and assessment of the mouse model for spinal cord injury. Zhongguo Xiu Fu Chong Jian Wai Ke Za Zhi. 2008 Aug;22(8):933-8. Chinese. PMID: 18773808.

25. Butler A, Hoffman P, Smibert P, Papalexi E, Satija R. Integrating single-cell transcriptomic data across different conditions, technologies, and species. Nat Biotechnol. 2018 Jun;36(5):411–420. doi: 10.1038/nbt.4096. Epub 2018 Apr 2. PMID: 29608179; PMCID: PMC6700744.

26. Macosko EZ, Basu A, Satija R, Nemesh J, Shekhar K, Goldman M, Tirosh I, Bialas AR, Kamitaki N, Martersteck EM, Trombetta JJ, Weitz DA, Sanes JR, Shalek AK, Regev A, McCarroll SA. Highly Parallel Genome-wide Expression Profiling of Individual Cells Using Nanoliter Droplets. Cell. 2015 May 21;161(5):1202–1214. doi: 10.1016/j.cell.2015.05.002. PMID: 26000488; PMCID: PMC4481139.

27. Aran D, Looney AP, Liu L, Wu E, Fong V, Hsu A, Chak S, Naikawadi RP, Wolters PJ, Abate AR, Butte AJ, Bhattacharya M. Reference-based analysis of lung single-cell sequencing reveals a transitional profibrotic macrophage. Nat Immunol. 2019 Feb;20(2):163–172. doi: 10.1038/s41590-018-0276-y. Epub 2019 Jan 14. PMID: 30643263; PMCID: PMC6340744.

28. Sullivan M P, Torres S J, Mehta S, et al. Heterotopic ossification after central nervous system trauma: a current review. Bone & joint research, 2013, 2(3): 51–57.

29. Mabbott NA, Baillie JK, Brown H, Freeman TC, Hume DA. An expression atlas of human primary cells: inference of gene function from coexpression networks. BMC Genomics. 2013 Sep 20;14:632. doi: 10.1186/1471-2164-14-632. PMID: 24053356; PMCID: PMC3849585.

30. Oyinbo CA. Secondary injury mechanisms in traumatic spinal cord injury: a nugget of this multiply cascade. Acta Neurobiol Exp (Wars). 2011;71(2):281–99. PMID: 21731081.

31. Couillard-Despres S, Bieler L, Vogl M. Pathophysiology of traumatic spinal cord injury[J]. Neurological aspects of spinal cord injury, 2017: 503–528.

32. Anwar MA, Al Shehabi TS, Eid AH. Inflammogenesis of Secondary Spinal Cord Injury. Front Cell Neurosci. 2016 Apr 13;10:98. doi: 10.3389/fncel.2016.00098. PMID: 27147970; PMCID: PMC4829593.

33. Orr MB, Gensel JC. Spinal Cord Injury Scarring and Inflammation: Therapies Targeting Glial and Inflammatory Responses. Neurotherapeutics. 2018 Jul;15(3):541–553. doi: 10.1007/s13311-018-0631-6. PMID: 29717413; PMCID: PMC6095779.

34. Jones TB, McDaniel EE, Popovich PG. Inflammatory-mediated injury and repair in the traumatically injured spinal cord. Curr Pharm Des. 2005;11(10):1223–36. doi: 10.2174/1381612053507468. PMID: 15853679.

35. Pineau I, Sun L, Bastien D, Lacroix S. Astrocytes initiate inflammation in the injured mouse spinal cord by promoting the entry of neutrophils and inflammatory monocytes in an IL-1 receptor/MyD88-dependent fashion. Brain Behav Immun. 2010 May;24(4):540–53. doi: 10.1016/j.bbi.2009.11.007. Epub 2009 Nov 22. PMID: 19932745.

36. Elkabes S, DiCicco-Bloom EM, Black IB. Brain microglia/macrophages express neurotrophins that selectively regulate microglial proliferation and function. J Neurosci. 1996 Apr 15;16(8):2508–21. doi: 10.1523/JNEUROSCI.16-08-02508.1996. PMID: 8786427; PMCID: PMC6578768.

37. Chamak B, Morandi V, Mallat M. Brain macrophages stimulate neurite growth and regeneration by secreting thrombospondin. J Neurosci Res. 1994 Jun 1;38(2):221–33. doi: 10.1002/jnr.490380213. PMID: 8078107.

38. Dusart I, Schwab ME. Secondary cell death and the inflammatory reaction after dorsal hemisection of the rat spinal cord. Eur J Neurosci. 1994 May 1;6(5):712–24. doi: 10.1111/j.1460-9568.1994.tb00983.x. PMID: 8075816.

39. Neirinckx V, Coste C, Franzen R, Gothot A, Rogister B, Wislet S. Neutrophil contribution to spinal cord injury and repair. J Neuroinflammation. 2014 Aug 28;11:150. doi: 10.1186/s12974-014-0150-2. PMID: 25163400; PMCID: PMC4174328.

40. Farooque M, Isaksson J, Olsson Y. Improved recovery after spinal cord trauma in ICAM-1 and P-selectin knockout mice. Neuroreport. 1999 Jan 18;10(1):131–4. doi: 10.1097/00001756-199901180-00024. PMID: 10094148.

41. Miron V E, Franklin R J M. Macrophages and CNS remyelination[J]. Journal of neurochemistry, 2014, 130(2): 165–171.

42. Niehaus JK, Taylor-Blake B, Loo L, Simon JM, Zylka MJ. Spinal macrophages resolve nociceptive hypersensitivity after peripheral injury. Neuron. 2021 Apr 21;109(8):1274-1282.e6. doi: 10.1016/j.neuron.2021.02.018. Epub 2021 Mar 4. PMID: 33667343; PMCID: PMC8068642.

43. Wahane S, Zhou X, Zhou X, Guo L, Friedl MS, Kluge M, Ramakrishnan A, Shen L, Friedel CC, Zhang B, Friedel RH, Zou H. Diversified transcriptional responses of myeloid and glial cells in spinal cord injury shaped by HDAC3 activity. Sci Adv. 2021 Feb 26;7(9):eabd8811. doi: 10.1126/sciadv.abd8811. PMID: 33637528; PMCID: PMC7909890.

44. Milich LM, Choi JS, Ryan C, Cerqueira SR, Benavides S, Yahn SL, Tsoulfas P, Lee JK. Single-cell analysis of the cellular heterogeneity and interactions in the injured mouse spinal cord. J Exp Med. 2021 Aug 2;218(8):e20210040. doi: 10.1084/jem.20210040. Epub 2021 Jun 16. PMID: 34132743; PMCID: PMC8212781.

45. Kigerl KA, Gensel JC, Ankeny DP, Alexander JK, Donnelly DJ, Popovich PG. Identification of two distinct macrophage subsets with divergent effects causing either neurotoxicity or regeneration in the injured mouse spinal cord. J Neurosci. 2009 Oct 28;29(43):13435–44. doi: 10.1523/JNEUROSCI.3257-09.2009. PMID: 19864556; PMCID: PMC2788152.

46. David S, Kroner A. Repertoire of microglial and macrophage responses after spinal cord injury. Nat Rev Neurosci. 2011 Jun 15;12(7):388–99. doi: 10.1038/nrn3053. PMID: 21673720.

47. David S, Greenhalgh AD, Kroner A. Macrophage and microglial plasticity in the injured spinal cord. Neuroscience. 2015 Oct 29;307:311–8. doi: 10.1016/j.neuroscience.2015.08.064. Epub 2015 Sep 2. PMID: 26342747.

48. Murray PJ, Allen JE, Biswas SK, Fisher EA, Gilroy DW, Goerdt S, Gordon S, Hamilton JA, Ivashkiv LB, Lawrence T, Locati M, Mantovani A, Martinez FO, Mege JL, Mosser DM, Natoli G, Saeij JP, Schultze JL, Shirey KA, Sica A, Suttles J, Udalova I, van Ginderachter JA, Vogel SN, Wynn TA. Macrophage activation and polarization: nomenclature and experimental guidelines. Immunity. 2014 Jul 17;41(1):14–20. doi: 10.1016/j.immuni.2014.06.008. PMID: 25035950; PMCID: PMC4123412.

49. Byrne AM, Bouchier-Hayes DJ, Harmey JH. Angiogenic and cell survival functions of vascular endothelial growth factor (VEGF). J Cell Mol Med. 2005;9(4):777–794. doi:10.1111/j.1582-4934.2005.tb00379.x

50. Storkebaum E, Lambrechts D, Carmeliet P. VEGF: once regarded as a specific angiogenic factor, now implicated in neuroprotection. Bioessays. 2004 Sep;26(9):943–54. doi: 10.1002/bies.20092. PMID: 15351965.

51. Mackenzie F, Ruhrberg C. Diverse roles for VEGF-A in the nervous system. Development. 2012 Apr;139(8):1371–80. doi: 10.1242/dev.072348. PMID: 22434866.

52. Sondell M, Lundborg G, Kanje M. Vascular endothelial growth factor stimulates Schwann cell invasion and neovascularization of acellular nerve grafts. Brain Res. 1999 Nov 6;846(2):219–28. doi: 10.1016/s0006-8993(99)02056-9. PMID: 10556639.

